# Fleeing is Believing: Adaptive behavior under social threat as an inference process

**DOI:** 10.64898/2026.02.17.706393

**Authors:** Hridai S Khurana, Valeria Mussetto, Cornelius T Gross, Rory J Bufacchi

## Abstract

Adverse social experiences profoundly alter animal behavior, yet the processes underlying context-appropriate behavior selection based on prior social interactions remain poorly understood. Existing models capture statistical patterns but are not framed in mechanistic frameworks that explain individual variability and predict behavioral outcomes.

We address this gap by modeling social defeat in mice as a partially observable Markov decision process (POMDP), implementing a heterarchical agent architecture - a structured network of interacting modules balancing exploration and exploitation.

Our model successfully reconstructs observed behavioral motifs (e.g., investigation, hesitation, and flights), fits different mouse phenotypes (e.g., susceptible vs. resilient), and mechanistically captures the impact of social defeat as a parameter shift in the animal’s internal generative model. The model reproduces effects of interventions like optogenetic stimulation, and generates testable predictions for future experiments.

The model’s modular architecture enables natural extension to other behavioral domains including foraging and multi-agent interactions, representing a foundational step toward interpretable models of mouse behavior. By capturing how adverse social experiences reshape decision-making at the computational level, this work offers potential clinical relevance for trauma and anxiety disorders in humans.

## 2. Introduction

Defensive behavior in animals emerges from a continuous decision-making process that balances the imperative to gather information against the risk of physical harm(1– 4). While modern ethological methods excel at cataloging behavioral structure, they often stop short of formally explaining the computational logic driving an animal’s choices. To understand how adverse experiences reshape behavior and make quantitative predictions, we need normative models that map hidden internal beliefs to observed actions.

Current computational approaches face challenges bridging algorithmic theory and biological implementation. Descriptive methods including high-resolution tracking (DeepLabCut(5), MoSeq(6,7), etc), statistical descriptions (Lévy walks, etc), and neural dynamical systems models(8) characterize what the system does but not why specific policies emerge. Normative frameworks (e.g., Bayes-Adaptive Markov Decision Process(9)) have formalized exploration as risk-sensitive trade-offs. However, these models operate at the level of discrete behavioral states that must be predefined and fitted from observed transitions, limiting their ability to generate testable predictions about the continuous dynamics of actual behavior. Can we build an interpretable model of ecological defensive behavior that generates recognizable behavioral patterns from first principles?

The social defeat paradigm(10) in rodents offers a tractable testbed. In this protocol [Fig. 1], defeat by an aggressive conspecific drives robust, quantifiable behavioral adjustments in subsequent social interactions, often shifting behavior from social exploration toward social avoidance (11,12). Such responses can be adaptive, promoting coping strategies that reduce risk during future encounters. Yet they may also hold clinical significance, as overly rigid coping patterns following social stress are implicated in disorders such as major depression and social anxiety(13–15).

**Figure 1.**
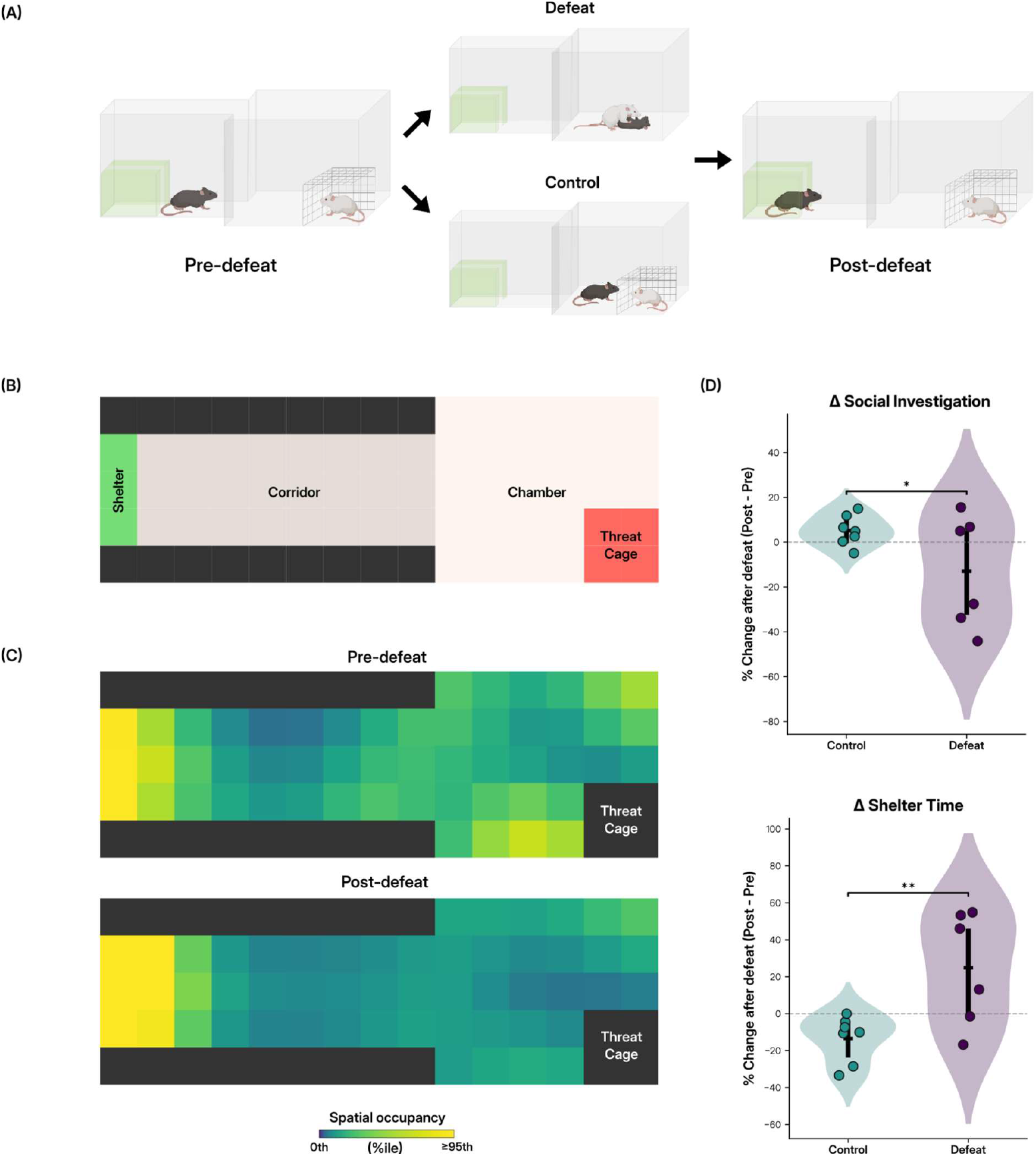
Social defeat experimental setup and empirical results (A) Experimental setup, showing pre-defeat investigation of a caged aggressive mouse (left), followed by defeat or control (middle), and post-defeat behavioral assessment (right). (B) Top-down view of the arena setup, divided into zones of the Shelter, Corridor, Chamber and Threat Cage. (C) Heatmaps showing log-scaled, smoothed spatial occupancy density (normalized to the 95th percentile) across the discretized arena pre- and post-defeat for an exemplary mouse. Following defeat, occupancy becomes concentrated in the shelter, with reduced exploration of the arena. (D) Percentage change between pre- and post-exposure phases, of time spent socially investigating and time spent in shelter, for the Control (teal) and Defeat (purple) groups. Individual data points represent individual mice; black markers indicate the group mean with error bars showing the 95% confidence interval of the mean. Positive values reflect an increase after the manipulation, whereas negative values reflect a decrease. The dashed horizontal line denotes zero change. Statistical significance was tested with unpaired t-test (**p < 0.01, *p < 0.05)

Critically, defeat does not produce a uniform phenotype: only a subset of individuals develop persistent avoidance, while others maintain social engagement. This variability presents a specific challenge: explaining how closely matched experiences can yield divergent behavioral outcomes.

We developed a heterarchical agent architecture implemented via Active Inference (16). Our model comprises three distinct, interacting modules: Spatio-Motor (executing navigation), Threat Identification (resolving stimulus identity), and Danger Context (encoding safety context) [Fig. 2(A)] - that collectively drive behavior by predicting sensory inputs, then adjusting actions based on prediction errors. This modular structure parallels the anatomical organization of defensive circuits as distinct computational subsystems. The question becomes whether such a model can fit real behavioral dynamics while offering a way to generate experimentally testable hypotheses about the neural computational mechanisms.

**Figure 2.**
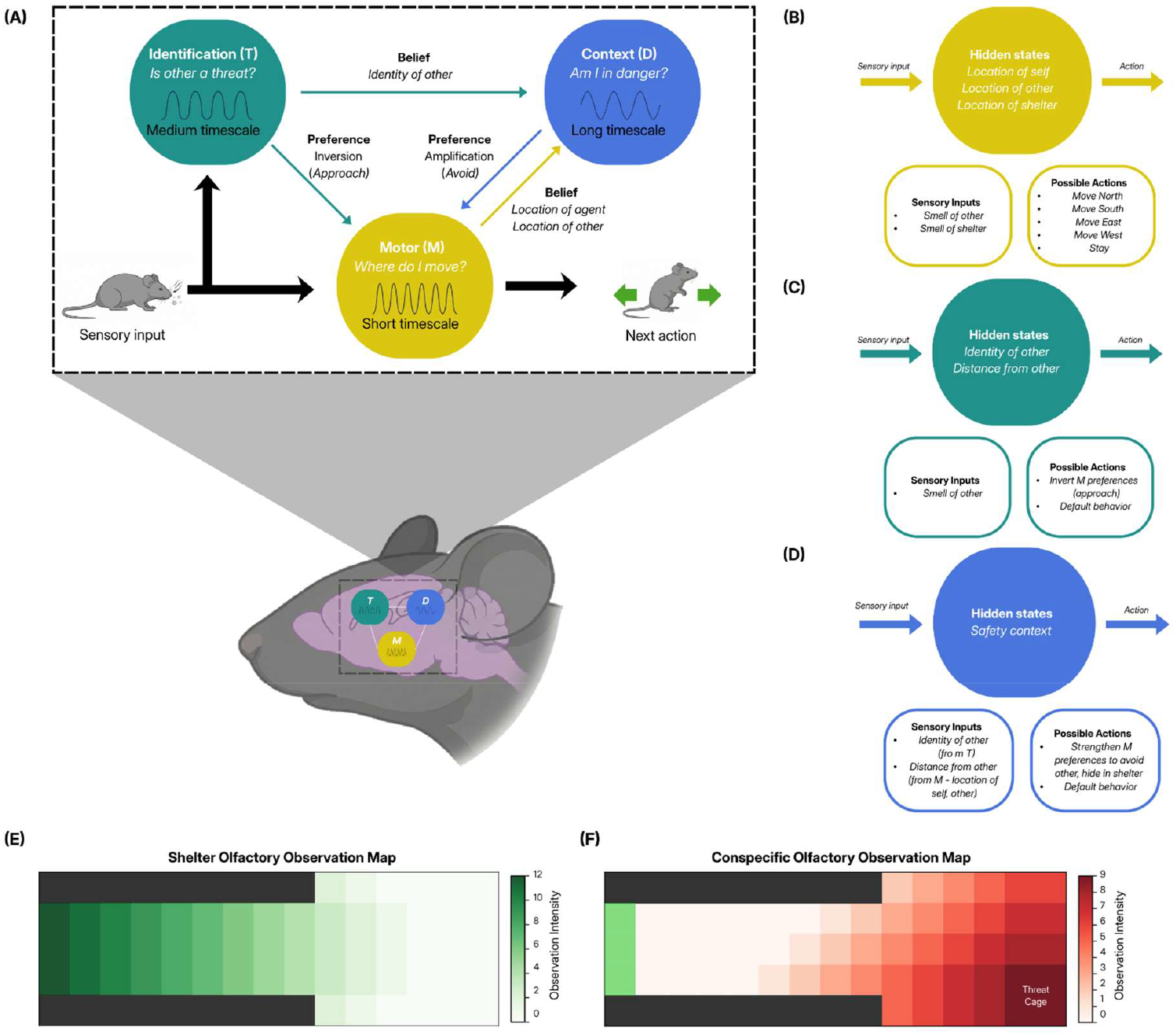
Model design (A) Schematic of interactions between modules of the heterarchical model. Sensory input is used to infer self-, other- and shelter-locations (by module M), and identity of the conspecific (by module T). Beliefs about threat and context modulate approach–avoidance preferences and shape subsequent actions. Module T takes action by biasing module M’s preferences to approach. These inferred locations and identity beliefs are used to determine the safety context (by module D), which biases preferences of module M towards escaping. (B-D) Factorized state–action mappings for (B) the Spatio-Motor module (M), (C) the Danger Context module (D), and (D) the Threat Identification module (T), showing sensory inputs, hidden states, and available actions. (E) Spatial map of sensory shelter observations received by the agent across arena locations. 13 discrete sensory observation states, peaking at the shelter location and decreasing with distance (F) Spatial map of conspecific sensory observations received by the agent across arena locations. 10 discrete observation states, peaking at the threat cage and decreasing with distance

By fitting the model to observed behavioral profiles from mice before and after social defeat, we find that phenotypic shifts can be captured by parameter updates within specific modules. The model produces continuous spatial trajectories and behavioral sequences (hesitation, flight, exploration) that emerge from belief updating rather than explicit programming. As a proof of concept for circuit-level prediction, we show that simulating optogenetic activation of the Danger Context module reproduces experimentally observed state-dependent escape responses elicited by optogenetic stimulation of the ventrolateral ventromedial hypothalamus (VMHvl)(12).

Our approach offers a starting point for linking behavioral dynamics to underlying computational processes. The modular architecture provides interpretability and potential extensibility to other behavioral domains, suggesting a path toward more general normative models of animal decision-making.

## 3. Results

### 3.1. Task Overview

We used a three days social defeat protocol adapted from Sureka et al(17) [Fig. 1(A)]. Each day, experimental mice were allowed to explore a two-compartment arena [Fig. 1(B)] consisting of a main chamber connected to a corridor terminating in a shelter, with an aggressive male mouse positioned in a wire-mesh cage in the chamber. During this period, the time spent investigating the aggressor as well as the time spent in the shelter were recorded. Next, the aggressor was released for a period sufficient to elicit social defeat (6-10 biting attacks) of the subject mouse. Control animals were run through the identical sequence, except that the aggressor remained confined behind the mesh barrier throughout [Fig. 1(A)]. On the last post-defeat test day, defeated mice displayed a robust avoidance phenotype, spending significantly less time near the caged aggressor and correspondingly more time in the shelter than control animals [Fig. 1(C), (D)].

### 3.2. Model Design

We developed a generative model that simulates defensive decision-making as a continuous active inference process. We first describe the model architecture and its generative dynamics independently of experimental condition. Differences between pre- and post-defeat behavior are introduced later in the Results (Section 2.5). The agent comprises three interacting modules [Fig. 2(A)]: Spatio-Motor [Fig. 2(B)], Threat Identification [Fig. 2(C)], and Danger Context [Fig. 2(D)]; each with distinct action timescales. Here, timescale refers to the typical rate at which a module accumulates evidence before updating its policy. The agent receives sensory olfactory^1^ inputs of shelter [Fig. 2(E)] and threat [Fig. 2(F)] from the environment.

Accordingly, the Spatio-Motor operates on a fast timescale to handle spatial navigation. The Threat Identification module operates on a slower cycle to resolve discrete uncertainty (e.g., ‘Threat’ vs. ‘Friendly’ conspecific), while the Danger Context module operates on the slowest timescale to encode persistent environmental states (‘Safety’ vs. ‘Danger’). However, these timescales are not rigid constraints, they can be triggered to update immediately when presented with unambiguous, high-precision evidence, allowing the system to react instantly to sudden threats.

These modules interact by exchanging inferred and preferred states. The system’s foundational survival drives are anchored in the Spatio-Motor module, which maintains a base preference for the shelter location and a strong aversion to observations of threat. The Danger Context module reinforces this by maintaining a persistent preference for ‘Safety’ over ‘Danger’ states.

In contrast, the Threat Identification module possesses no intrinsic preference for specific outcomes. Instead, it is driven purely by epistemic value—the imperative to resolve uncertainty. To minimize belief entropy regarding the intruder’s identity, this module adjusts the Spatio-Motor to temporarily invert its preferences. This interaction generates a curiosity-driven approach behavior that directly competes with the pragmatic drive for safety.

The model’s core dynamics emerge from the resolution of this conflict. To illustrate, we consider a typical behavioral episode (Fig. 3):

1. Investigation: The agent begins sheltered with high uncertainty [Fig. 3(A); time course up to the blue circle]. The Threat Identification module, prioritizing epistemic value of the conspecific [Fig. 3 (B),(C)], overrides the Spatio-Motor’s safety preferences, driving the agent to approach the caged mouse closer than it would under the Spatio-Motor module’s default shelter-seeking policy, to learn specifically about the caged mouse more effectively [Fig. 3(A); blue trajectory].
2. Identification: As the agent navigates closer, the precision of sensory observations increases [Fig. 3(A); blue trajectory to red circle]. Once the Threat Identification module resolves the target’s identity with a confidence exceeding the classification threshold [Fig. 3(B)], this belief is propagated to the Danger Context module.
3. Flight: Upon updating its belief to a ‘Danger’ context, the Contextual module releases the epistemic override and amplifies the Spatio-Motor’s base preference for the shelter. With uncertainty resolved beyond the threshold, and safety prioritized, the agent executes a ballistic flight back to the shelter [Fig. 3(A); time from red circle to end]. As the agent retreats and distance increases, sensory precision fades, causing the Threat Identification belief to rapidly decay back to baseline uncertainty [Fig. 3(B)]
4. Crucially, the Danger Context module bridges this sensory gap by maintaining the ‘Danger’ state, sustaining the ballistic flight even after the immediate evidence of threat has been lost [Fig. 3(C)]

**Figure 3.**
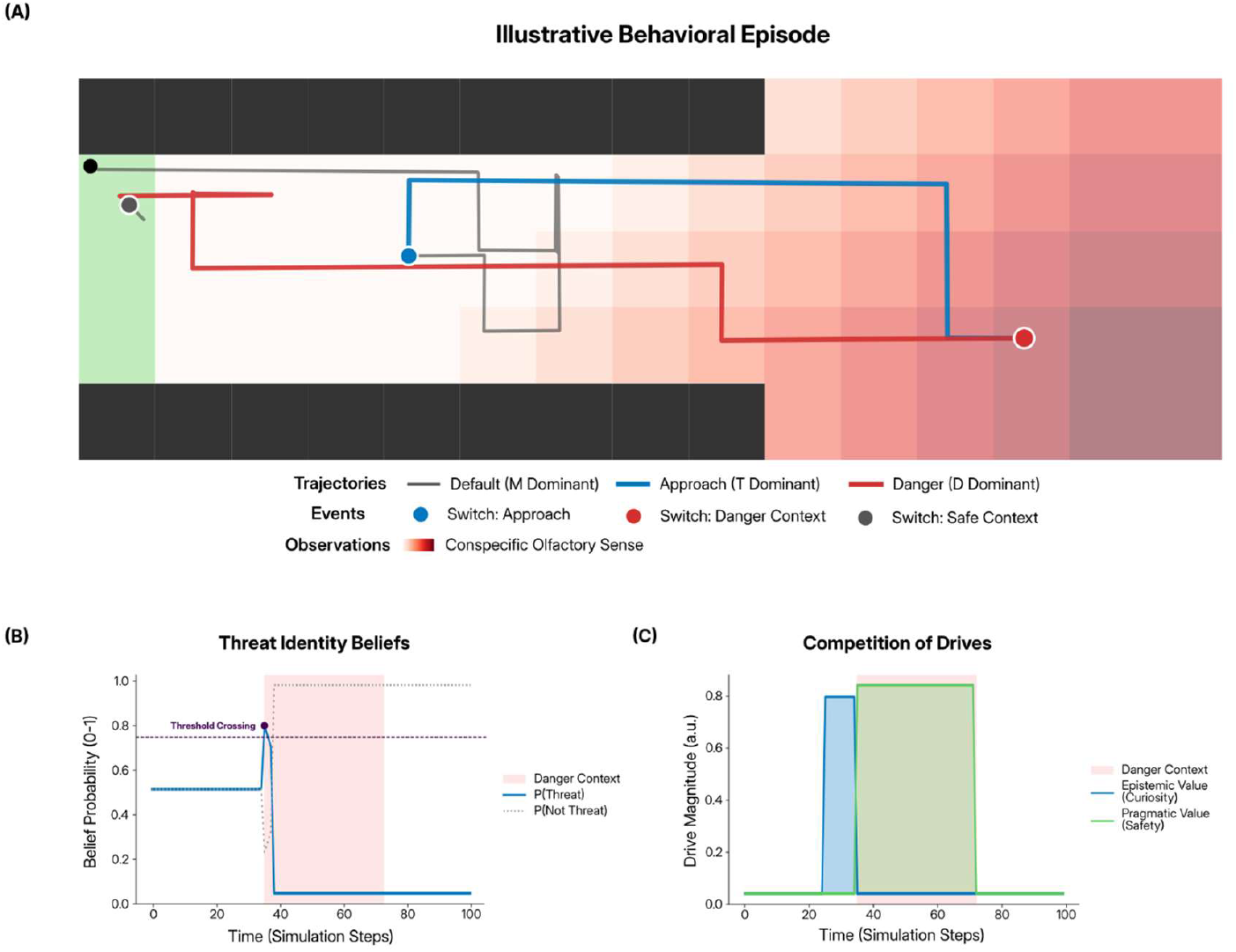
Illustrative behavioral episode arising from conflict resolution between epistemic and safety drives. (A) Example trajectory of the agent in the arena during a single episode. Red shading indicates the conspecific olfactory sensory intensity (darker is higher); green coloring indicates the shelter. The agent begins in the shelter under high uncertainty and initially approaches the caged conspecific (blue trajectory) as the Threat Identification module dominates the preferences of the Spatio-Motor module, and hence action selection to maximize epistemic value. The blue circle marks the onset of approach dominance; the red circle marks the point at which threat identity is resolved. After the belief that a threat is present is propagated to the Danger Context module, control shifts to danger context-driven avoidance, producing a ballistic return (flight) to the shelter (red trajectory). (B) Time course of threat identity beliefs. Increasing sensory precision during approach drives a rapid rise in the probability of the conspecific being a “threat,” which crosses a decision threshold and subsequently decays as the agent retreats and sensory evidence diminishes. (C) Competition between epistemic (curiosity) and pragmatic (safety) drives over time, with arbitrary units indicating the drive magnitude. Epistemic value dominates during investigation, but is taken over by pragmatic value after threat identification and inference of danger context, with sustained avoidance behavior and flight even after immediate sensory evidence of threat has weakened.

Note that while threshold-crossing triggers action from the Danger Context module, it can also transition to a danger state through its own temporal dynamics. Its slower timescale allows threat beliefs to accumulate even under ambiguous sensory evidence, enabling the model to simulate anxiety-like states that persist, and escapes that occur, without a clear relation to acute triggers.

### 3.3. Model captures the latent structure of empirical mouse behavior

We validated the model architecture by fitting model parameters to aggregate trajectories from experimental cohorts (refer to Methods Section 4.6) tracked via DeepLabCut(5). Using Bayesian optimization (with the Optuna toolbox(18), which uses Tree-Structured Parzen Estimator(19)), we minimized a composite loss function comparing simulated versus observed behavior across five aggregate metrics: shelter occupancy, social investigation duration, spatial transitions, exploratory diversity (spatial entropy), and immobility proportion. Due to time and computational limitations, a representative library of agents was generated and uniquely matched to empirical data of individual mice.

As a baseline for performance, we compared the fitted model against a stochastic null model (random walk) [Fig. 4(A,B)]. While the random walk failed to replicate the structured occupancy and transition patterns characteristic of the experimental subjects, the fitted model’s output closely recapitulated empirical distributions across all analyzed metrics. Even with the limitations of the matching process, the fitting yielded sub-sigma accuracy for the majority of the cohort, with a median of 0.56*σ*, indicating that the simulated agents are statistically indistinguishable from the experimental subjects (for more information, see Methods Section 4.6.6).

**Figure 4.**
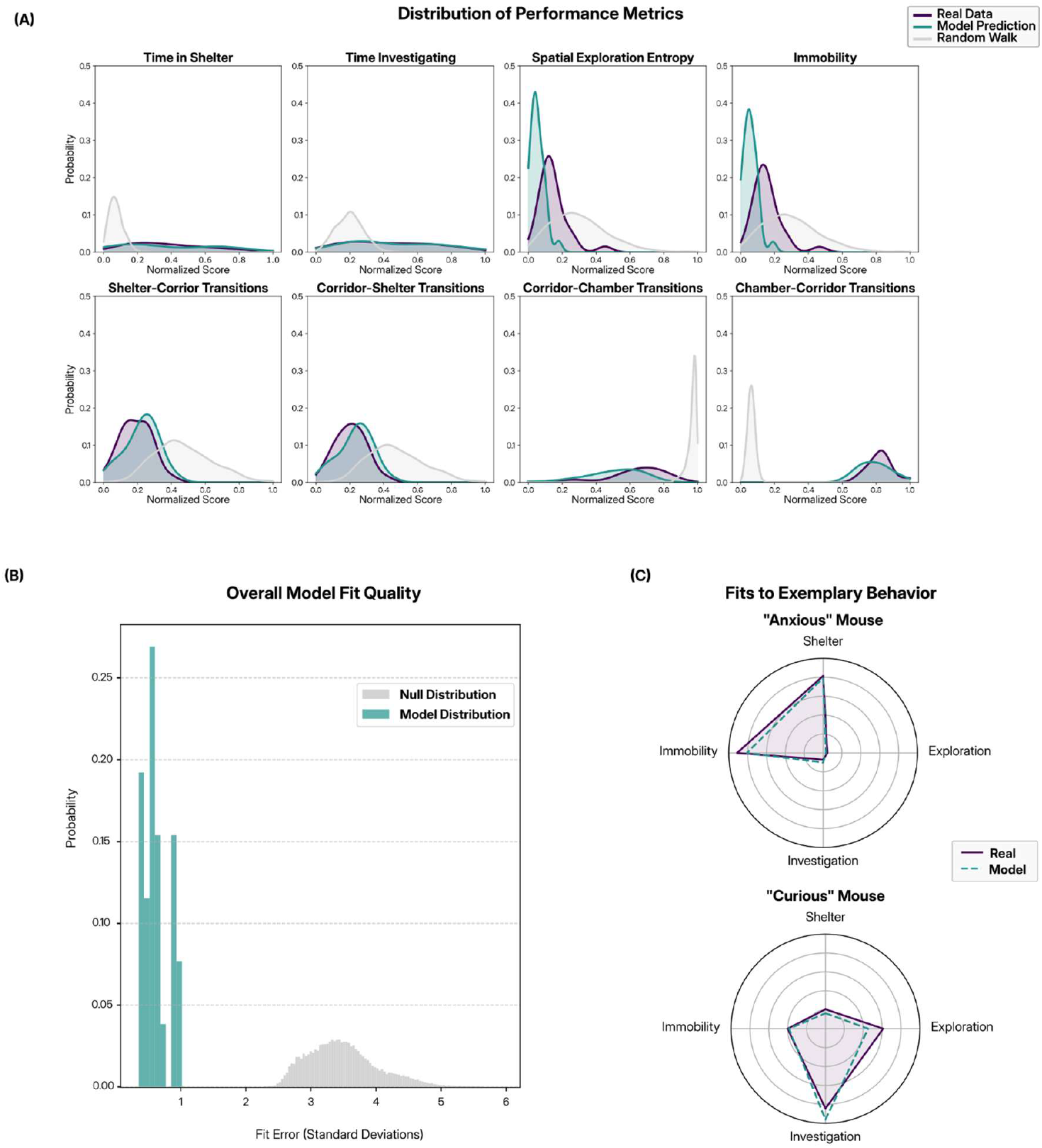
Model fit to empirical behavioral data. (A) Population-level distributions of normalized (min-max scaled over all distributions) behavioral metrics comparing real data (purple) with model predictions (teal); a random-walk baseline (gray) is shown for reference. Metrics include time allocation (shelter, investigation, immobility), spatial exploration entropy, and transition probabilities between arena zones (shelter, corridor, chamber). Across metrics, the model closely matches the empirical distributions and substantially outperforms the null baseline. (B) Overall model fit quality quantified by aggregate fit error (in standard deviations) across sessions. The model distribution is concentrated near low error values relative to the null distribution, indicating good global agreement with data, with median error of 0.56*σ* and max error of ∼1*σ*. (C) Radar plots illustrating fits to exemplary behavioral phenotypes on either extreme of the spectrum (“anxious” and “curious” mice), showing close correspondence between real (solid) and model (dashed) profiles across key behavioral dimensions.

### 3.4. Behavioral regimes emerge from the heterarchical model

The heterarchical model captures a broad spectrum of mouse individual variability by generating distinct behavioral regimes. These regimes are not pre-programmed but emergent, allowing the model to fit individual differences across a population.

#### 3.4.1. Behavioral Phenotypes

We demonstrate the model’s flexibility by fitting it to exemplary behavioral phenotypes from mice at two extremes of the behavioral spectrum, with other individuals falling between these poles:

- **The “Curious” Phenotype:** Exhibits wide-ranging exploration and full-arena coverage
- **The “Anxious” Phenotype:** Displays high shelter-dependency, with investigations rarely extending beyond the corridor

The model’s fit to example episodes of such mice at the extremes of the behavioral spectrum are shown in Fig. 4(C).

#### 3.4.2. Behavioral Motifs (Sequential Actions)

Within these overarching regimes, the model reconstructs recognizable, species-typical sequences. These motifs occur dynamically based on the agent’s internal state:

- **Hesitation:** Approaching the chamber followed by a localized retreat
- **Flight:** Rapid, ballistic trajectories from the stimulus area directly back to the shelter

The model’s ability to replicate these dynamics (Fig. 5) demonstrates that both stable individual traits and reactive defensive sequences can emerge from the proposed framework, without requiring separate hand-coded rules.

**Figure 5.**
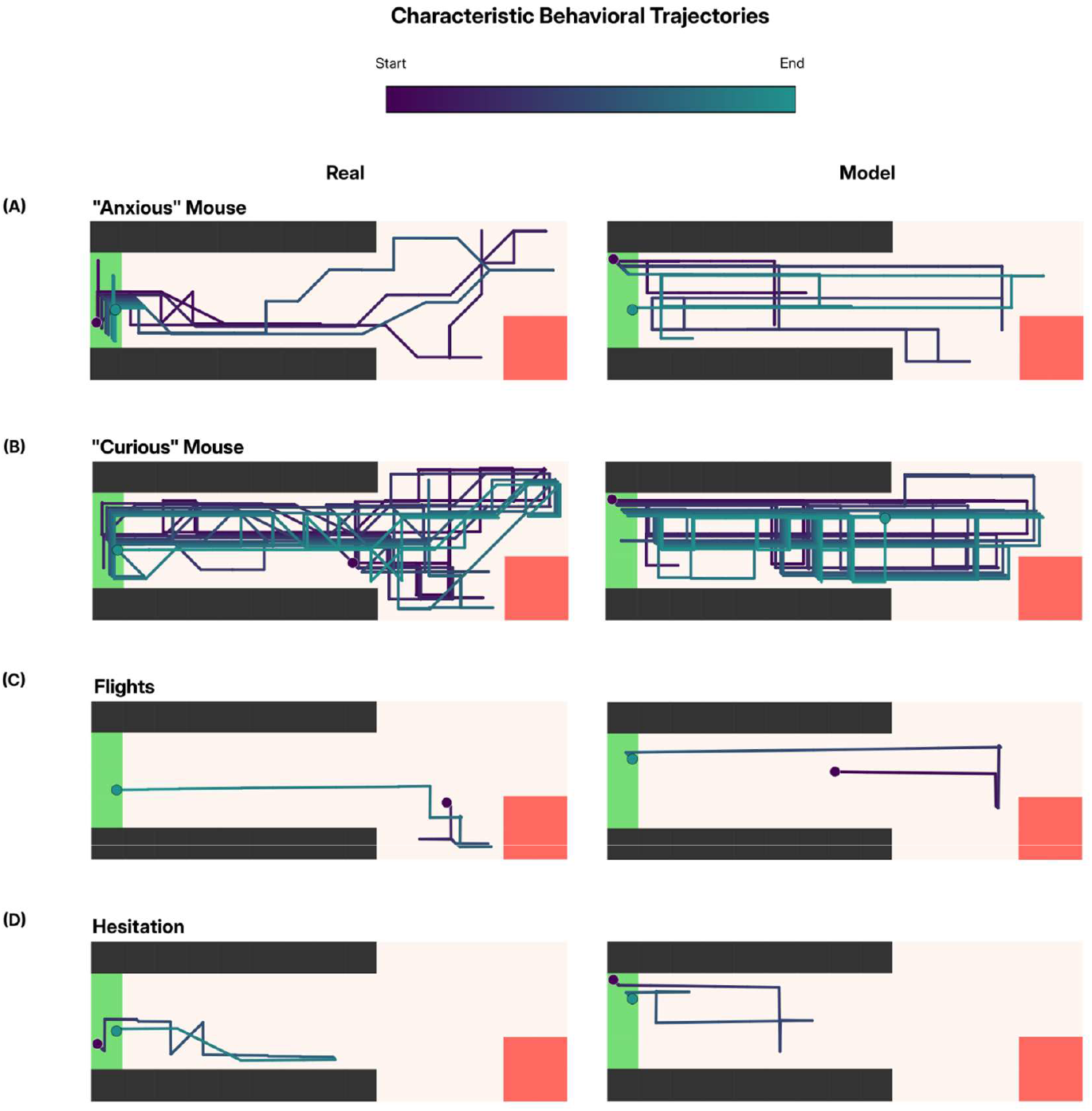
Characteristic behavioral trajectories reproduced by the model. Example trajectories from empirical data (left) and corresponding model simulations (right) for representative behavioral phenotypes and motifs. Trajectories are colored by time from start (purple) to end (teal). Across cases, the model qualitatively reproduces spatial structure and behavioral motifs observed in real trajectories, without fitting the model to the actual trajectories themselves. (A) Full experiment (2000 timesteps, equivalent to 5 mins of empirical recording) trajectory of an “anxious” mouse, exhibiting high time in shelter engagement and limited exploration. (B) Full experiment (2000 timesteps, equivalent to 5 mins of empirical recording) trajectory of a “curious” mouse, showing extensive exploration around the arena with frequent transitions. (C) Short trajectory snippet (∼200 timesteps, equivalent to ∼30 secs of empirical recording) showing flight behaviors characterized by rapid, ballistic movement across the arena, from investigating the threat cage to the shelter. (D) Short trajectory snippet (∼200 timesteps, equivalent to ∼30 secs of empirical recording) showing hesitation behaviors, marked by short excursions out of the shelter followed by retreat, without going out into the chamber.

### 3.5. Social defeat induces a parametric shift within the model

To quantify the computational impact of social defeat, we optimized the model parameters separately to fit the observed behavior of individual mice before and after their social encounters. By projecting these fits into a reduced parameter space [Fig. 6(A)], we can visualize how the underlying decision-making variables change for each animal.

**Figure 6.**
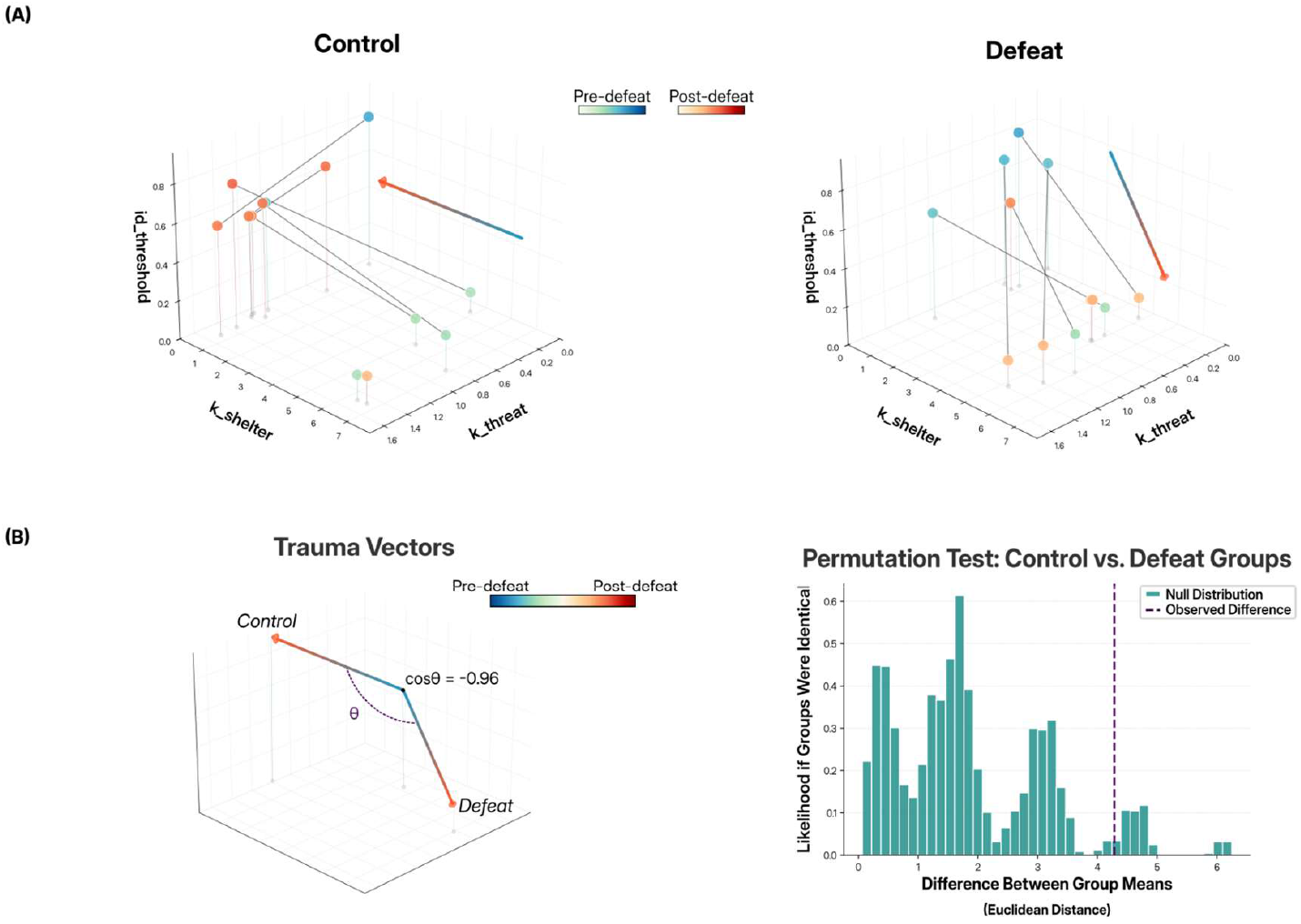
Social defeat induces a shift in model parameter space (A) Projections of selected fit model parameters (threshold for identity detection, preference for shelter, preference for threat - refer to Methods Section 4.6.5) for control (left) and defeat (right) mice, shown pre- (cool colors) and post- (warm colors) defeat. Each grey line connects pre-to post-session fits for a single mouse, illustrating within-subject changes in decision-related parameters. Arrows show the net “Trauma vector” pre-to post-defeat parameter change in each condition. Defeated mice exhibit a directional displacement in parameter space following defeat, relative to the controls. (A) *Left:* “Trauma vectors” summarizing the net pre–post parameter change for each group reveal opposing directions between control and defeat conditions (cos θ = − 0.96). *Right:* Permutation test comparing group mean vectors (Euclidean distance) shows separation between control and defeat trajectories relative to a null distribution. Although underpowered (p=0.069) given the sample size, the observed effect size (Mahalanobis *d* = 1.49) indicates a biologically meaningful divergence, supporting the interpretation that social defeat is captured by stable, systematic updates to a small set of internal model parameters.

In the defeat group, the model reveals a consistent, directional shift in parameter space following the aggressive encounter. While the starting points vary due to individual baseline differences, the net shift vectors for most defeated mice start in the same region of the parameter space and move in a similar direction, suggesting a common response to the experience is captured by the model.

In contrast, the control group exhibits a different pattern of movement within the parameter space. While these mice also show shifts between sessions, the net direction of these changes quantified by their “Trauma Vectors” are distinct from those observed in the defeat group (cosθ = -0.96, permutation test, p=0.069, Mahalanobis d=1.49) [Fig. 6(B)]. Given the small sample size (n=7 and 6 for control and defeat groups respectively), the substantial Mahalanobis distance suggests a biologically relevant divergence in behavioral trajectories that may be underpowered at the standard alpha level (0.05). small set of internal model parameters. These shifts reflect differences between parameters fitted independently to pre- and post-defeat behavior, rather than online parameter updates within the agent. The behavioral consequences of social defeat can therefore be summarized as systematic changes in a low-dimensional set of inferred model parameters.

### 3.6. Simulated optogenetics on model can recreate findings on dynamic encoding of social threat in VMHvl

Next, we simulated the effects of optogenetic VMHvl stimulation. This analysis serves three purposes. First, it tests the model’s ability to capture the neural circuit-level adaptations observed after social defeat as changes in latent belief. Second, it illustrates how the model’s predictions can constrain the space of candidate hypotheses. Third, it shows how the model can inform the design of future experiments that discriminate between these candidates.

Previous experiments(12) demonstrate that this stimulation triggers robust escape in defeated mice but not in naive controls, indicating a state-dependent gating mechanism. We hypothesize the Danger Context module parallels the VMH in function, and modeled optogenetic intervention as an exogenous bias on this module toward a *Danger* state. In contrast to social defeat, which is represented here as an offline difference in fitted parameters, optogenetic stimulation is modeled as an explicit online perturbation to the model.

While multiple hypotheses could plausibly account for defeat-induced sensitization, we focus here on a small set to illustrate both the predictive utility and the falsifiability of the model. We tested three hypotheses:

1. **Sensitization to danger (“Model S”):** Defeat sensitizes the Danger module to input from the Threat module. It increases the spatial range over which social stimuli are treated as potentially dangerous, effectively lowering the distance threshold at which “*Danger*” evidence is accumulated.
2. **Biased Identity Priors (“Model T”):** Defeat increases the prior probability that the social target is a threat (hyper-vigilance in Threat Identification).
3. **Biased Context Priors (“Model D”):** Defeat increases the prior probability that the environment itself is unsafe (persistent risk in Danger Context).

Under baseline pre-defeat parameters, *in silico* optogenetic stimulation failed to elicit flight in any model, mirroring biological controls [Fig. 7(A)]. After applying parameter shifts optimized to fit the statistical distribution of empirical post-defeat behavior, both Model T and Model D reproduced the elevated flight rates observed in defeated mice under optogenetic stimulation [Fig. 7(B)].

**Figure 7.**
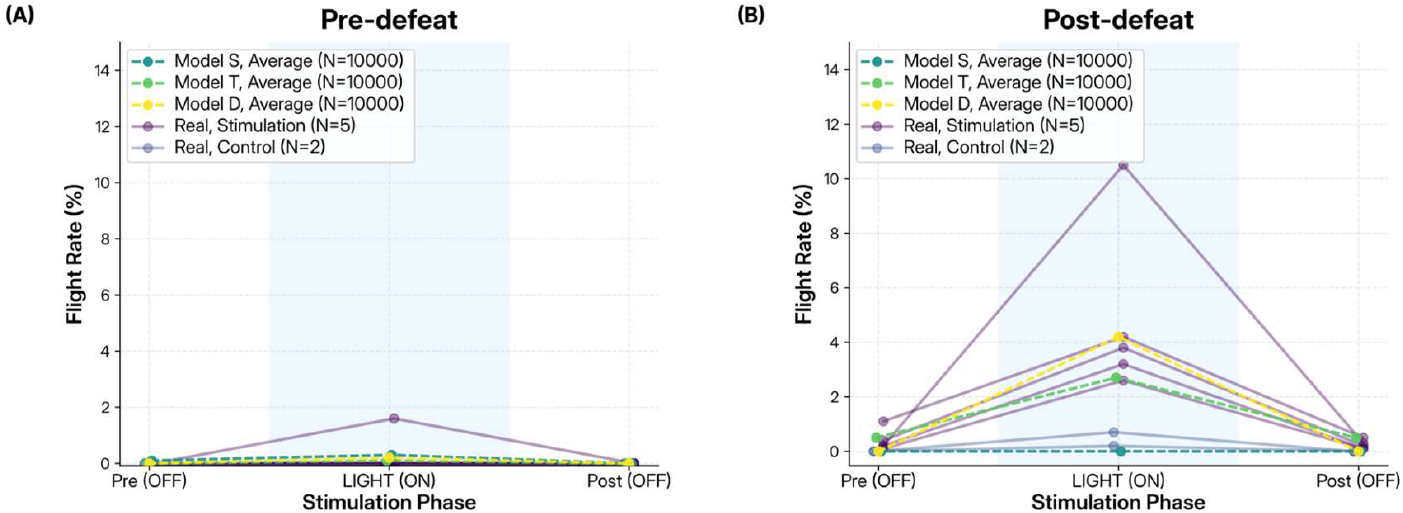
State-dependent optogenetic stimulation effects emerge from illustrative hypothesized defeat-induced belief changes. Empirical data was taken from Fig. 2(e, f) in Krzywkowski et al., 2020(12).Together, these results show that the model can capture and possibly explain state-dependent efficacy of VMHvl stimulation, illustrated by: either biasing of social threat identity priors or a bias toward a danger context. (A) *Pre-defeat*. Flight rates in response to optogenetic VMHvl stimulation for real mice and model simulations under baseline (pre-defeat) parameters. In both Model T (biased identity priors) and Model D (biased context priors), exogenous activation of the Danger Context module fails to elicit escape, mirroring the lack of stimulation-evoked flight observed in empirical data. (B) *Post-defeat*. After applying the defeat-induced prior biases optimized to match empirical behavior, simulated optogenetic stimulation produces robust increases in flight rate in both models, closely matching the responses observed in defeated mice.

Model S, by contrast, did not alter the model’s optogenetic response under the conditions tested, in both pre- and post-defeat states [Fig. 7(A),(B)]. Although expanded threat generalization is a behaviorally plausible consequence of defeat, in the experimental configuration used here the relevant social interactions occur at distances where this manipulation does not change the inferred state. As a result, increasing the spatial range of danger sensitivity alone produces no effect on the efficacy of VMHvl stimulation.

These results demonstrate a form of constrained computational degeneracy: multiple hypotheses can account for the defeat state-dependent impact of VMHvl stimulation, but of those tested, only modified social identity or environmental danger priors (models T and D) are expressed in these conditions. In the Discussion Section 3.3, we discuss how to experimentally differentiate between these two hypotheses.

## 4. Discussion

### 4.1. A Computational Account of Social Defeat

Social defeat is associated with divergent behavioral outcomes following similar aggressive encounters, ranging from social avoidance to preserved social engagement. Our model reproduces these behaviors [Fig. 5] and represents individual differences as variation within its parameter space [Fig. 6(A)], while remaining interpretable by mapping behavior onto cognitive parameters like inferred danger, current assessment of threat, and preferences for sensory stimuli.

This degree of interpretable variability distinguishes the approach from existing models, which either characterize what the system does without explaining why specific policies emerge(5–8), or explain the policies without characterizing inter-individual variability and low-level behavioural dynamics (e.g. movement)(9).

The model also shows potential to capture how social defeat reorganizes decision-making [Fig. 6]. Relative to controls, defeated animals show a directional shift in parameter space, characterized by a lower threshold for committing to threat-related inferences of social cues consistent with increased anxiety(20,21), and a stronger bias toward shelter-seeking behavior(22). The difference in directionality and substantial Mahalanobis distance (d = 1.49) support biological relevance, though larger cohorts are needed for robustness.

### 4.2. Neural Implementation

The modular architecture suggests putative circuit correspondences(23). The output of the Spatio-Motor module could align with evolutionarily conserved structures in the brainstem, such as the periacqueductal gray (PAG), required for the execution of instinctive defensive motor patterns(23–25). The Danger Context module may parallel hypothalamic, in particular VMH, involvement in encoding threat imminence and internal defensive state(11,24,26), consistent with defeat-induced potentiation of VMH responses and gating of escape behavior(12). The Threat Identification module could correspond to the medial amygdala (MeA), given its role in processing socially relevant chemosensory cues and differentiating threat-related from neutral stimuli(27,28).

If this mapping is approximately correct, the model predicts that selectively disrupting Medial or Posterior Amygdala (MeA/PA) projections to VMH would cause impaired social threat identification(29), while leaving defensive response execution intact. In this scenario, defensive motor programs would remain intact but would be less reliably or appropriately engaged due to degraded upstream threat signaling, rather than a failure of downstream motor execution. This interpretation remains tentative, as both the proposed circuit correspondences and the division of labor between identification, contextual modulation, and execution within these regions are incompletely resolved. Further work is required to determine which defensive responses would be preserved, the conditions under which they would be triggered, and how such manipulations would affect responses to non-olfactory cues and non-social threats, or whether parallel amygdalo–hypothalamic or cortical pathways could compensate for loss of MeA input. Addressing these issues will be essential for determining the specificity and limits of the proposed mapping.

### 4.3. Experimental Predictions for Neurostimulation

To illustrate the model’s capability of capturing the neural plasticity observed in social defeat, we first narrowed a broader hypothesis space to mechanisms that reproduce the observed state-dependent effects of optogenetic VMH stimulation. In doing so, we excluded the hypothesis that the danger-assessing module is sensitised to input from the threat-assessment module after social defeat.

We were left with two defeat-induced modification hypotheses: biased identity priors (“Model T”) and biased contextual priors (“Model D”). We then examined how these distinct defeat-induced modifications account for the experimental results observed in defeated animals(12). Both hypotheses reproduce the defensive behaviors observed experimentally in the presence of subordinate conspecifics.

Therefore, our model demonstrates that the two hypotheses are currently computationally degenerate under the conditions tested. As an illustration of the model’s predictive utility, we propose a test to distinguish between them: VMH stimulation in defeated mice in the absence of a conspecific altogether. Model D predicts escape behavior, driven by activation of a generalized danger state, whereas Model T predicts little or no response, as no ambiguous social stimulus is available for classification.

The outcome of this experiment would help reveal whether defeat reorganizes defensive behavior at the level of social inference or through a global shift in threat state. Although alternative explanations may exist, here we deliberately focus on these two mechanistically distinct hypotheses as a minimal and illustrative test case.

### 4.4. Limitations and Future Directions

The model omits several ethologically important phenomena of defensive behaviour: thigmotaxis(30), social buffering(31,32), context-dependent fear responses(33), habituation(34), and broader defensive repertoires including stretch-attend postures(35,36); to name a few. Behavior is modeled through centroid trajectories only, omitting postural and kinematic features. Many such features could be incorporated through model extensions like additional state factors and expanded observations.

While the present model offers a viable explanation for defensive behaviour, alternative hypotheses can currently not be ruled out for several reasons. First, we evaluate limited model variants. Broader exploration of alternative architectures, learning rules, and parameter ranges is needed for model identifiability in future work. Second, the framework has so far been applied to a single paradigm, without incorporating concurrent neural recordings, limiting validation of the proposed circuit mappings. Even within the social defeat paradigm, the modest sample size (n = 13) and marginal group significance (p = 0.069) underscore the need for larger cohorts to confirm computational signatures of defeat. Finally, although the current parameter optimization fits the empirical data, it admits multiple behaviorally equivalent solutions, which precludes circuit-level inference. In principle, more rigorous, module-specific parameter updates could instead generate testable hypotheses about the neural circuits involved in defeat-induced plasticity.

Future work should also incorporate neural recordings to validate belief state mappings, expand to richer behavioral measures, and test the framework across multiple paradigms and species. For example, the modeling principles could apply to individual differences in trauma response, underlying vulnerability and resilience to post-traumatic psychopathology(37). By representing defeat-induced changes as shifts in interpretable computational parameters, this framework may offer a formal language for describing such differences. While highly speculative, this raises the possibility that analogous parameters could eventually be estimated from human behavior.

### 4.5. Broader Applicability

The modular architecture allows extension to other behavioral domains (e.g., foraging(38); social interaction via theory-of-mind–like inference) with relatively limited modification, and could serve as a foundation for a general model of mouse behavior. Such a model would enable systematic generation of testable hypotheses, support principled comparison across experimental paradigms, and reduce the need for exhaustive empirical exploration by narrowing the space of plausible behavioral mechanisms.

Normative and active inference–based generative models using the principles demonstrated here have also been applied to other naturalistic behaviors in rodents and other systems like ant colonies, illustrating the broader utility of this class of models(39–41). The modular architecture demonstrated in this work could help such models generalize further to describe more kinds of ecological behaviors, while still maintaining interpretability.

### 4.6. Conclusion

This work demonstrates that normative models can capture naturalistic defensive behavior while maintaining interpretability. By framing effects of social defeat as a parameter shift to an animal’s internal generative model, we unify behavioral dynamics, individual variability, and circuit-level hypotheses. The modular architecture shows how shared principles can generate diverse defensive behaviors. It provides a foundation for studying how adverse experience reshapes the computations underlying social decision-making, and for future extensions generalizing the framework across tasks.

## 5. Methods

### 5.1. Active Inference Framework

Each of the modules in the model is in itself an agent using Active Inference, a normative framework which posits that agents act to minimize the expected free energy of their future observations(42–44). Formally, the agent operates within a Partially Observable Markov Decision Process (POMDP) governed by an internal generative model. This model is parameterized by a likelihood matrix *A*, which maps hidden states to sensory observations (*P*(*o* | *s*)), a transition matrix *B*, which defines the probabilistic evolution of states given a policy *P*(*s*_*t*+1_ | *s*_*t*_, π), and a preference distribution *C*, which encodes the agent’s prior beliefs about preferred outcomes *P*(*o*).

The behavioral loop proceeds in two distinct phases: perception and planning. In the perception phase, the agent infers the current hidden state of the environment *s*_*t*_ from sensory observations *o*_*t*_. This is achieved by inverting the likelihood model *A* to minimize the Variational Free Energy *F*.

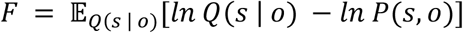

This minimization approximates the posterior belief *Q*(*s* | *o*) that best explains the sensory data, effectively converting raw sensation into a probabilistic understanding of the current context.

In the planning phase, the agent projects this belief into the future to select a policy *π* (a sequence of actions). Using the transition matrix *B*, the agent predicts future states and evaluates them against its preferences *C*. It does this by selecting the policy that minimizes the Expected Free Energy *G*(*π*_*t*_) evaluated over future time steps *t* ∈ [1, *N*], where N denotes the planning horizon - the number of time points over which the agent can anticipate future states. This decomposes into two distinct components:

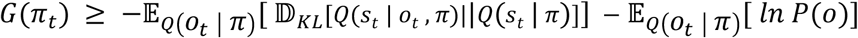

The first term, Epistemic Value, computes the expected information gain (Bayesian surprise) between the prior and posterior, representing the entropy of the predicted observations, driving the agent toward states that resolve uncertainty (e.g., exploration). The second term, Pragmatic Value, represents the expected log probability of observing the preferred observation under the selected policy, intuitively computing how likely it is that this policy will drive the agent toward goal-directed behavior (e.g., safety or reward). By minimizing this dual objective, the agent naturally alternates between exploratory and defensive behaviors depending on its current uncertainty and environmental context. We used the PyMDP Python library to implement this framework(45).

### 5.2. Model Environment

The environment was modeled as a 2-dimensional discrete grid world [Fig. 1(B)] comprising 57 distinct spatial states. The arena dimensions were made to match the geometry of the physical experimental arena used in the behavioral assay. The arena contained two distinct features:

- Shelter zone, comprising three grid states on the end of the corridor.
- Conspecific cage, comprising four grid states on one corner of the chamber.

The environment generated sensory observations based on the agent’s spatial proximity to these features, which can be interpreted as simulating a mix of multiple sensory modalities:

1. **Conspecific Signal:** The conspecific source (cage) emitted a position-dependent signal. The intensity of this signal was determined by the Manhattan distance (since the agent moves only in the cardinal directions) between the agent and the cage location.
2. **Shelter Signal:** Similarly, the shelter emitted a location signal, providing a “homing” gradient, enabling the agent to navigate toward safety.
3. **Agent Location:** The agent received feedback regarding its current grid coordinate, which was used to update its internal belief about its position within the arena.

At each timestep, the environment calculated the agent’s new position based on its chosen motor action, enforced boundary constraints (if trying to enter walls or the cage, the agent does not change position), and updated the sensory signals accordingly. The environment was deterministic in its physical dynamics.

### 5.3. Generative Model

The agent’s internal model is defined by a Partially Observable Markov Decision Process (POMDP) structured into tuples of matrices (*A, B, C, D, E*). We distribute these matrices across three distinct, interacting modules: **Spatio-Motor (M), Threat Identification (T)**, and **Danger Context (D)**.

1. **Spatio-Motor Module (M)** This module handles low-level navigation and spatial inference.
  - **State Space (*S***_***M***_): Each state is one of the 57 spatial locations in the discretized arena.
  - **Observations (*O***_***M***_): The agent receives three sensory channels: Self-location, Conspecific Sensory Signal (scalar 0-9), and Shelter Sensory Signal (scalar 0-12).
  - **Actions (*U***_***M***_): 5 possible moves {*Up, Down, left, Right, Stay*}.
  - **Likelihood (*A***_***M***_): Maps hidden states (spatial locations) to observations. Proprioception is deterministic, while Conspecific/Shelter signals are probabilistic, with precision decaying over distance.
  - **Transitions (*B***_***M***_): Encodes physical movement physics. Maps the effect of actions *U*_M_ ∈ {*Up, Down, left, Right, Stay*} to transition the agent to adjacent cells. Actions are restricted to cardinal directions to limit computational cost and avoid diagonal trajectories that yield non-physical speed advantages.
  - **Preferences (*C***_***M***_): Defines the agent’s immediate goals over observations, which are to stay near shelter (positive utility) and avoid threats (negative utility). Crucially, this matrix is dynamic; it is modulated by higher-level modules to switch the agent’s drive between exploring (investigating the threat) and fleeing (seeking shelter).
  - **Priors (*D***_***M***_): Prior belief on state, initialized as a uniform probability over the state space, to simulate the agent not immediately knowing the locations of itself, shelter, or threat.
  - **Habits (*E***_***M***_): Encodes strong priors over actions. Here, the *Stay* action receives a bias, to simulate energy cost or immobility observed.
2. **Threat Identification Module (T)** This module acts as the identification system, inferring the identity of ambiguous social targets.
  - **State Space (*S***_***T***_): Two hidden state factors: *Identity* {*Not threat, threat*} and *Proximity* [0,9], where distances beyond 9 are considered equal.
  - **Observations (*O***_***T***_): Receives the sensory input for Conspecific Sensory Signal (scalar 0-9).
  - **Action (*U***_***T***_): The module selects between {*Null, Approach*} actions. If the *Approach* action is selected, the T module inverts the Motor module’s preferences to assign a negative utility to the shelter and positive to threat observations, facilitating investigation.
  - **Likelihood (*A***_***T***_): Maps hidden Identity and Proximity states to observations. The precision of this mapping is distance-dependent, governed by a fitted parameter *sensory_slope* (see Methods Section 4.6.4); signals are noisy at a distance and precise up close.
  - **Transitions (*B***_***T***_): Identity is stable (a threat does not spontaneously become safe). Proximity states update based on the Motor module’s movement.
  - **Preferences (*C***_***T***_): No utility to observations, agent does not prefer a specific Conspecific Sensory Signal.
  - **Priors (*D***_***T***_): Prior belief on state, initialized as a uniform probability over identity and distance factors, to simulate the agent not immediately knowing the current state.
  - **Habits (*E***_***T***_): No bias on any action.
3. **Danger Context Module (D)** This module encodes whether the agent is in danger or not.
  - **State Space (*S***_***D***_): Two context hidden states: {*Safe, Danger*}
  - **Observations (*O***_***D***_): A joint signal derived from the *T* module’s identity belief {*Not threat, threat*}, and *distance from the conspecific* [0, 9] derived from the *M* module.
  - **Action (*U***_***D***_): Chooses between {*Null, Avoid*}. When the *Avoid* action is chosen, this module strengthens the Motor module’s default preferences (*C*_*M*_) with a high negative utility for the Conspecific proximity observation and a high positive utility for the shelter, resulting in a flight response.
  - **Likelihood (*A***_***D***_): Maps context state to observations. Observations where the identity belief is of *threat* and distance is close enough (modulated by a distance threshold parameter) are mapped to *Danger* state context, others are *Safe*.
  - **Transitions (*B***_***D***_): *Avoid* action switches context to *Safe*.
  - **Preferences (*C***_***D***_): Negative utility to observations that map to *Danger* context.
  - **Priors (*D***_***D***_): Initialized as a uniform probability over context state.
  - **Habits (*E***_***D***_): No bias on any action.

### 5.4. Simulation Protocol

The simulation is structured as an active inference process across the three levels, shown in [Algorithm 1]. At each time step *t*, each module in the agent performs state estimation (inference, minimizing variational free energy). The M module performs action selection (minimizing expected free energy) at each time step, while higher levels (T and D) perform action selection every *ticks*_*T*_ or *ticks*_*D*_ time steps, or with event-based triggers, upon reaching specific belief thresholds to modulate the agent’s goal preferences (represented by the *C* vector).

To simulate long-term pre-existing beliefs, we implement a forgetting mechanism where state beliefs (posteriors) decay toward their generative prior at rates *η*_*M*_, *η*_*T*_, *η*_*D*_ after each step. As a result, high-precision beliefs are maintained only through ongoing interaction with the environment, rather than defaulting to generalized assumptions about longer-term state.

#### Algorithm 1

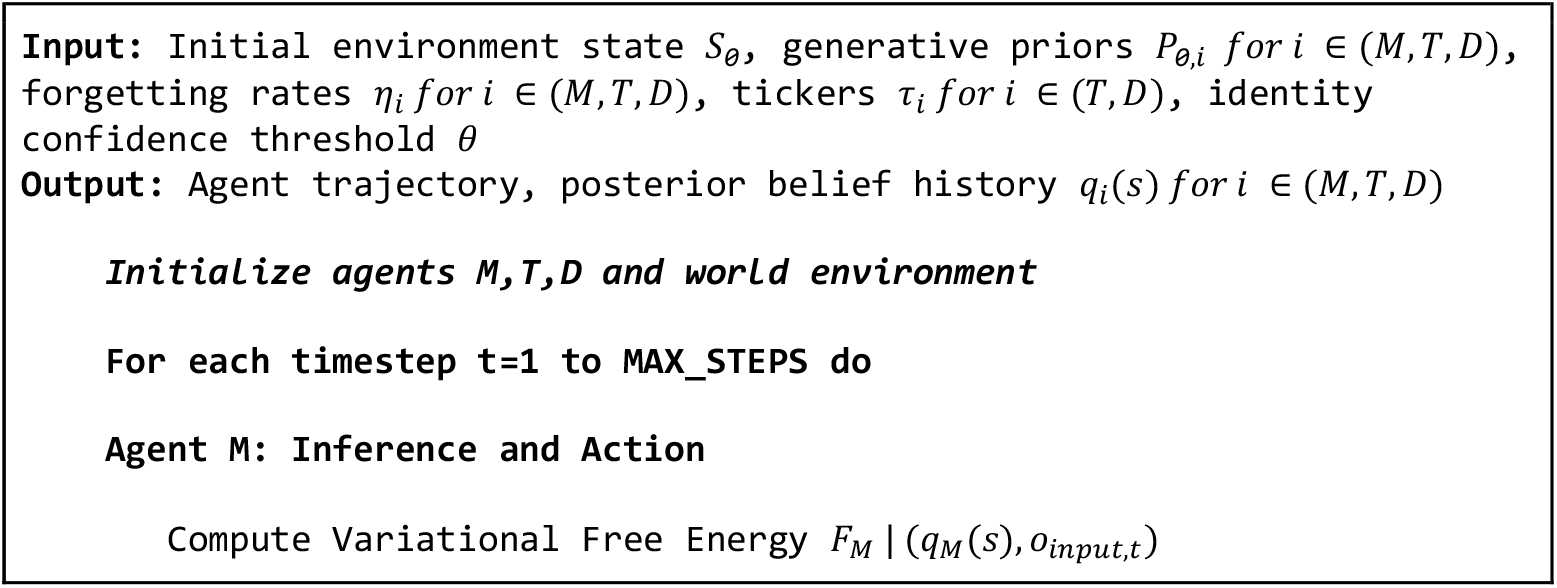
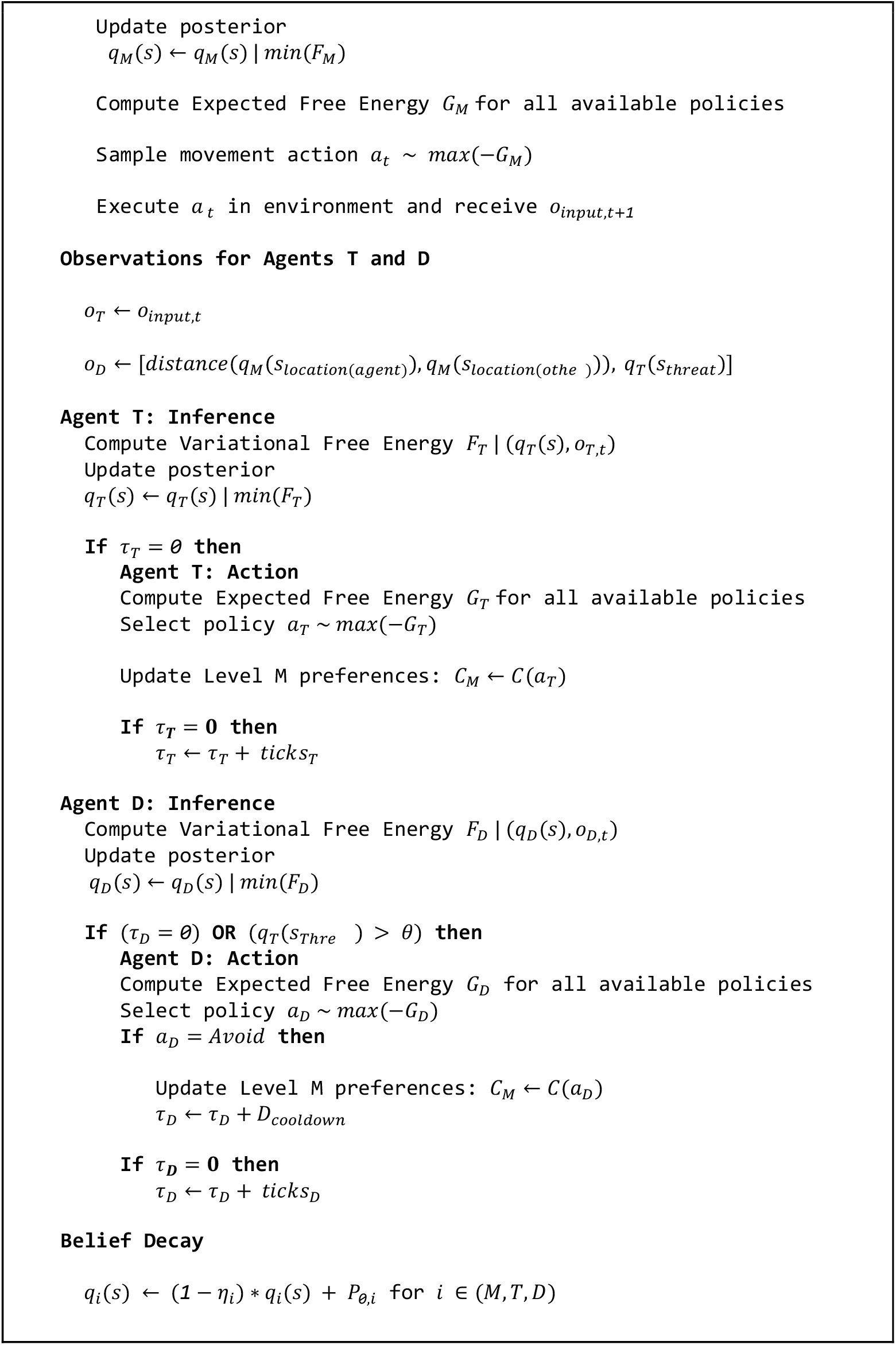
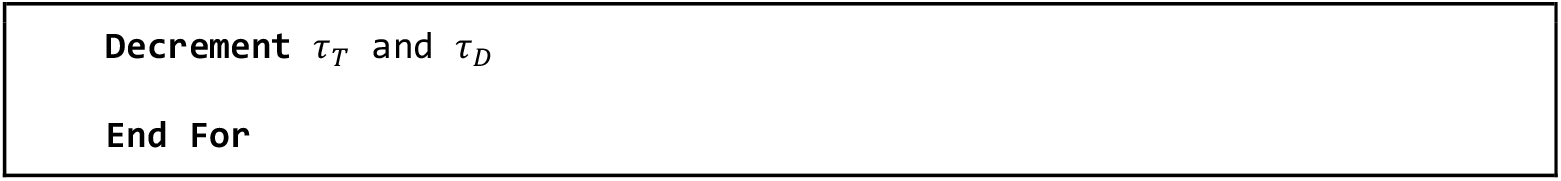

### 5.5. Social Defeat Experiments

#### 5.5.1. Animals

All experimental procedures involving the use of animals were carried out in accordance with EU Directive 2010/63/EU and under the approval of the EMBL Animal Use Committee and the Italian Ministry of Health License 183/2024-PR to Cornelius Gross. C57BL/6 J wild-type mice were obtained from local EMBL colonies. Adults (>8 weeks old) were used in all experimental procedures. The experiments were performed in male mice, since the social defeat procedure relies on interactions with aggressive males. Aggressor mice were single housed CD1 adult retired male breeders purchased from Charles Rivers Laboratories. CD1 mice were pre-screened for aggressiveness(46). CD1 mice were considered aggressive if they defeated different C57BL/6J males across three consecutive resident–intruder sessions performed in the C57BL/6J home cage. A defeat was scored when the CD1 delivered 4-6 attack bouts that elicited defensive responses in the C57BL/6J mouse (escape, freezing, upright postures). All animals were maintained in a temperature and humidity-controlled environment with 12 hr light and 12 hr dark cycle, and food and water were provided ad libitum.

#### 5.5.2. Behavioral testing

The social defeat procedure was adapted from (Sureka et al., 2025)(17). The apparatus consisted of a plexiglas chamber (25 × 25 × 25 cm) connected via a 12-cm-wide doorway with a sliding door to a plexiglas corridor (34 × 12 × 25 cm). A rectangular shelter with a narrow entrance (8 × 12 x 7 cm; red Plexiglas) was placed at the distal end of the corridor, and a wire-mesh cage was positioned in the corner opposite the corridor opening inside the larger chamber. Experimental C57BL/6J mice were single-housed for a week before testing. They were then habituated to the empty apparatus for 5 min on two consecutive days. Subsequently, mice underwent three sessions of habituation to the aggressor (habituation days 1–3) consisting of four phases: (i) exploration of the empty apparatus for 5 min; (ii) investigation, during which an aggressive CD1 mouse was placed inside the wire-mesh cage in the chamber while the experimental mouse freely explored the apparatus for 5 min; (iii) restriction, during which the experimental mouse was confined to the chamber for 5 min by closing the sliding door while the CD1 remained confined within the wire-mesh cage; and (iv) post-restriction exploration of the entire apparatus for 5 min after reopening the sliding door. Next, on defeat days 1 and 2, the same sequence was performed for the No Defeat group. For the Defeat group, during the restriction phase the CD1 mouse was released from the wire-mesh cage into the chamber to permit a social defeat encounter, which was terminated after 6 attacks or 10 min (whichever occurred first). Immediately thereafter, the CD1 mouse was returned to the wire-mesh cage and the experimental mouse completed the post-restriction phase (5 min free exploration of the full apparatus). On the final day (post-defeat test day), all mice underwent the same procedure until the investigation phase only. To prevent familiarity effects, each experimental mouse was exposed to a different CD1 aggressor on each day, and each CD1 mouse was used for no more than one defeat session per day. The apparatus was cleaned between animals by wiping sequentially with ethanol and water.

All behavioral sessions were video-recorded using a Basler camera positioned above the apparatus (top-down view) at 25 frames per second. Pose was estimated using DeepLabCut(5) with a ResNet-50 backbone. A network was trained on manually labeled frames sampled from a subset of videos spanning sessions and subsequently applied to all videos to extract x–y coordinates for eight body landmarks. Outliers in the tracked data were subsequently identified and corrected using SimBA software(47).

### 5.6. Biological Validation and Parameter Optimization

To ensure the model captures realistic ethological dynamics, we fit the agent’s parameters to biological data derived from the social defeat experiment.

#### 5.6.1. Experimental Dataset

We utilized a dataset of mouse trajectories extracted via DeepLabCut (5)from experimental videos, consisting of 13 mice split into Control (n=7) and Defeat (n=6) groups. These continuous trajectories were mapped into the 57-state grid matching the aspect ratio of the arena, and subsampled to 2000 time steps, corresponding to approximately 5 minutes of real-time investigation phase of the behavioral experiment. This discretization preserves the key spatial topology - shelter, corridor, and threat cage, allowing for direct comparison between biological and simulated agents. The investigation phase just before the first defeat was taken as the pre-defeat behavior, and investigation after the final defeat day was taken as post-defeat behavior for both Control and Defeat groups.

#### 5.6.2. Optimization Framework

We employed Optuna(18), a Bayesian optimization framework, to tune the agent’s free parameters. Specifically, we used the Tree-Structured Parzen Estimator (TPE)(19) sampler to explore the high-dimensional parameter space efficiently. The optimization targeted five key internal parameters that govern the agent’s cognitive and motor dynamics:

1. *id_threshold*: The confidence level required by the Threat Identification (T) module to classify a signal as a threat.
2. *sensory_slope*: The rate at which sensory precision decays with distance, determining the agent’s sensory acuity or uncertainty about distal objects.
3. *k_shelter*: The magnitude of the preference (utility) for the shelter.
4. *k_threat*: The magnitude of the preference (utility) for avoidance of the threat, calibrating the agent’s risk/reward sensitivity.
5. *bias_stay*: The transition probability bias (habit) assigned to the “Stay” action in the Motor module, serving as a proxy for motor lethargy or decision latency.

#### 5.6.3. Ethological Metrics and Loss Function

The optimization objective was to minimize a composite loss function representing the divergence between the simulated agent’s behavior and the “Average Mouse” profile (derived from the control group). We defined a set of eight ethological metrics to capture distinct facets of defensive behavior:

- **Time in Shelter:** The fraction of total time steps the agent occupies the designated shelter cells.
- **Social Investigation Time:** The fraction of time spent in the interaction zone (cells within distance ≤2 of the threat), conditioned on temporal continuity (occupying the zone for at least 3 consecutive frames) to filter out transient crossings.
- **Zone Transitions:** The frequency of border crossings between the three functional zones (Shelter ↔ Corridor, Corridor ↔ Chamber), capturing the dynamism of the agent’s behavior.
- **Spatial Exploration Entropy:** The Shannon entropy of the occupancy distribution over the 57 states, quantifying the diversity of exploration.
- **Immobility:** The proportion of time steps where the agent’s location remains unchanged (*location*_*t*_ = *location*_*t*-1_), measuring general motor inactivity.

The global loss *L* was calculated as a weighted sum of the squared z-scores for each metric *k*, normalizing the simulation output *s*_*k*_ against the biological mean *μ*_*k*_ and standard deviation *σ*_*k*_ of the target cohort:

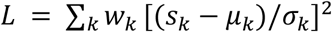

where *w*_*k*_ represents the importance weight of each metric. We prioritized the primary ethological readouts - **Time in Shelter** and **Investigation Time**, with higher weights (w=2.0), while all other metrics like transitions and entropy were weighted lower (w=0.5), for a total weight of 7.5. This hierarchical weighting ensures the optimizer prioritizes ethological validity (where the mouse spends its time) while still constraining the agent to realistic motor patterns (how it moves).

#### 5.6.4. Parameter Estimation via Representative Profile Matching

To model individual variability without the computational cost of independent optimization for every subject, we employed a profile-matching strategy. We first conducted optimization runs across a manifold of representative parameter configurations to generate a library of behavioral profiles. Each experimental subject was then uniquely mapped to the parameter set from this library that minimized its specific loss function.

We assessed the quality of these matches by converting the fitting loss into standard deviations relative to the experimental data variance (*σ* = *L*/∑_*k*_ *w*_*k*_).

This metric confirms that the discrete parameter library provided sufficient resolution to capture individual differences with high fidelity.

The matching process yielded sub-sigma accuracy for the majority of the cohort. The median subject was matched with a precision of 0.56*σ*, and even the outlier cases remained close to 1.0*σ*. This indicates that the simulated agents are statistically indistinguishable from the experimental subjects, as the model deviation falls well within the natural behavioral variance of the biological population.

### 5.7. *In-silico* Optogenetic Stimulation

To simulate optogenetic activation of the ventromedial hypothalamus (VMH), we introduced an exogenous bias to the Danger Context module agent’s belief state at the moment of conspecific encounter.

Stimulation was modeled as an additive boost to the posterior belief in the *Danger* state, applied immediately after sensory inference but prior to action selection. When the agent crossed the distance threshold, the belief state was modulated according to:

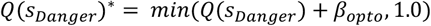

Where *Q*(*s*_*Danger*_) represents the belief of the Danger Context module derived from sensory evidence and *β*_*opto*_ is the stimulation intensity.

## 6. Code and Data Availability

All custom code used for modeling and analysis is available on GitHub at https://github.com/hridaik/social-defeat

## 7. Acknowledgements

We thank Flàvia Ferrús Marimón, Leiron Ferrarese, and Sofija Perovic for their helpful comments and suggestions on earlier drafts of this manuscript.

## Footnotes

1 Although framed as *olfactory* sensory inputs, these inputs are not restricted to literal smell. Rather, *olfactory* serves as an intuitive proxy for any sensory signal that informs shelter or threat detection. Accordingly, *olfactory* and *sensory* are used interchangeably throughout the paper.

